# An evolutionary portrait of the progenitor SARS-CoV-2 and its dominant offshoots in COVID-19 pandemic

**DOI:** 10.1101/2020.09.24.311845

**Authors:** Sudhir Kumar, Qiqing Tao, Steven Weaver, Maxwell Sanderford, Marcos A. Caraballo-Ortiz, Sudip Sharma, Sergei L. K. Pond, Sayaka Miura

## Abstract

We report the likely most recent common ancestor of SARS-CoV-2 – the coronavirus that causes COVID-19. This progenitor SARS-CoV-2 genome was recovered through a novel application and advancement of computational methods initially developed to reconstruct the mutational history of tumor cells in a patient. The progenitor differs from the earliest coronaviruses sampled in China by three variants, implying that none of the earliest patients represent the index case or gave rise to all the human infections. However, multiple coronavirus infections in China and the USA harbored the progenitor genetic fingerprint in January 2020 and later, suggesting that the progenitor was spreading worldwide as soon as weeks after the first reported cases of COVID-19. Mutations of the progenitor and its offshoots have produced many dominant coronavirus strains, which have spread episodically over time. Fingerprinting based on common mutations reveals that the same coronavirus lineage has dominated North America for most of the pandemic. There have been multiple replacements of predominant coronavirus strains in Europe and Asia and the continued presence of multiple high-frequency strains in Asia and North America. We provide a continually updating dashboard of global evolution and spatiotemporal trends of SARS-CoV-2 spread (http://sars2evo.datamonkey.org/).

## Main

Despite an unprecedented scope of global genome sequencing of Severe acute respiratory syndrome coronavirus 2 (SARS-CoV-2) and a multitude of phylogenetic analyses^1–5^, the early evolutionary history of SARS-CoV-2 remains unclear. Sophisticated investigations have found that traditional molecular phylogenetic analyses do not produce reliable evolutionary inferences about the early history of SARS-CoV-2 due to low sequence divergence, a limited number of phylogenetically informative sites, and the ubiquity of sequencing errors^6–8^. In particular, the root of the SARS-CoV-2 phylogeny remains elusive^9,10^ because the closely-related non-human coronavirus (outgroups) more than 1,100 base differences from human SARS-CoV-2 genomes, as compared to fewer than 30 differences between human SARS-CoV-2 genomes’ sequenced early on (December 2019 and January 2020)^7,9–15^. Without a reliable root of the SARS-CoV-2 phylogeny, one cannot accurately reconstruct the most recent ancestor sequence. Consequently, we cannot determine if any of the coronaviruses isolated to date carried the genome of the most recent common ancestor (progenitor) of all human SARS-CoV-2 infections. Knowing the progenitor genome will help us determine how close the earliest patients sampled in China represent are to “patient zero,” i.e., the first case of human transmission.

The orientation and order of early mutations giving rise to common coronavirus variants will be misled if the earliest coronavirus isolates are incorrectly used to root the SARS-CoV-2 phylogenies^3,16–18^. The earliest investigations of COVID-19 patients and their coronaviruses’ genomes already reported the presence of multiple variants^19,20^, and genomes of viral samples from December 2019 had as many as five differences from each other. These observations require an explicit test of the assumption that one of the early sampled coronavirus genomes was the most recent common ancestor (progenitor) of all the strains infecting humans. Traditionally, the ancestral sequence of organisms is estimated by using a rooted phylogeny^21,22^. This ancestral sequence can then be compared with sequenced genomes to find the one that is most similar to that of the inferred progenitor and/or placed closest to the root in the phylogeny. However, as noted above, attempts using *ad hoc* and traditional methods are fraught with difficulties and have produced contradictory results^9,10^. Some methods also incorporate sampling times in phylogenetic inference, but they favor placing the earliest sampled genomes at or near the root of the tree^10^. This practice introduces a degree of circularity in testing the hypothesis that the earliest sampled genomes were ancestral because sampling time is used in the inference procedure.

## Results and Discussion

### A mutational order approach for SARS-CoV-2

We applied a mutation order approach (MOA) that directly reconstructs the ancestral sequence and the mutational history of genomes^23–25^ without inferring a phylogeny as an intermediate step. MOA is often used to reconstruct the evolutionary history of tumor cells that evolve clonally and without recombination. This approach is well-suited for analyzing SARS-CoV-2 genomes because of their quasi-species evolutionary behavior (clonal) and because of the lack of evidence of significant recombination within human outbreaks, both of which preserve the collinearity of variants in genomes. This feature permits effective use of shared co-occurrence of variants in genomes, as well as the frequencies of individual variants, to infer mutational history, notwithstanding the presence of sequencing errors and other artifacts^23,26^ (see *Methods*). We advanced MOA for application in the analysis of SARS-CoV-2 genomes because the normal cell sequence in tumors provides a direct way to establish the ancestral (non-cancerous) genome. Such a direct ancestor is not available for coronaviruses in which the closest outgroup sequences are over 30-times more different than any two human strains. We also devised a bootstrap approach to place confidence limits on the inferred mutation order in which bootstrap replicate datasets are generated by sampling genomes with replacement (see *Methods*).

We analyzed two snapshots of the fast-growing collection of SARS-CoV-2 genomes to make inferences and assess the robustness of the inferred mutational histories to the growing genome collection, expanding at an unprecedented rate. The first snapshot was retrieved from GISAID^27^ on July 7, 2020, and consisted of 60,332 genomes. Of these, 29,681 were selected because they were longer than the 28,000 bases threshold imposed (29KG dataset) and did not include an excessive number of unresolved bases in any genomic regions. This second snapshot was acquired on October 12, 2020, from GISAID and contained 133,741 genomes, of which 68,057 genomes met the inclusion criteria (68KG dataset).

In the following, we first present results from the 29KG dataset and then evaluate the concordance of the mutational history inferred by using an expanded 68KG dataset, which establishes that the conclusions are robust to the sampling of genomes. We then applied mutational fingerprints inferred using the 68KG dataset to an expanded dataset of 172,480 genomes (sampled on December 30, 2020; 172KG) to track global spatiotemporal dynamics SARS-CoV-2. We have also set up a live dashboard showing regularly updated results because the processes of data analysis, manuscript preparation, and peer-review of scientific articles are much slower than the pace of expansion of SARS-CoV-2 genome collection. Also, we provide a simple “in-the-browser” tool to classify any SARS-CoV-2 genome based on key mutations derived by the MOA analysis (http://sars2evo.datamonkey.org/).

### Mutational history and progenitor of SARS-COV-2

We used MOA to reconstruct the history of mutations that gave rise to 49 common single nucleotide variants (SNVs) in the 29KG dataset (**Fig. 1**). These variants occur with greater than 1% variant frequency (*vf* > 1%; **Fig. 2a**). For ease of reference, we used the inferred mutation history to denote key groups of mutations by assigning Greek symbols (μ, ν, α, β, γ, δ, and ε) to them. Individual mutations were assigned numbers and letters based on the reconstructed order and their parent-offspring relationships (**Extended Data Table 1**). We estimated the timing of mutation for each mutation based on the timestamp of the viral samples’ genome sequences in which it first appeared (**Extended Data Table 1**, see *Methods*). The inferred mutation order generally agreed with the temporal pattern of the first appearance of variants in the 29KG dataset. The sampling time of 47 out of 49 mutations was greater than or equal to the first appearance of the corresponding preceding mutation in mutational history. The exceptions were seen only for two low-frequency offshoot mutations (β_3b_ and β_3c_; see *Methods*). This concordance provides independent validation of the reconstructed mutation graph because neither sampling dates nor locations were used in MOA analysis.

**Fig. 1.**
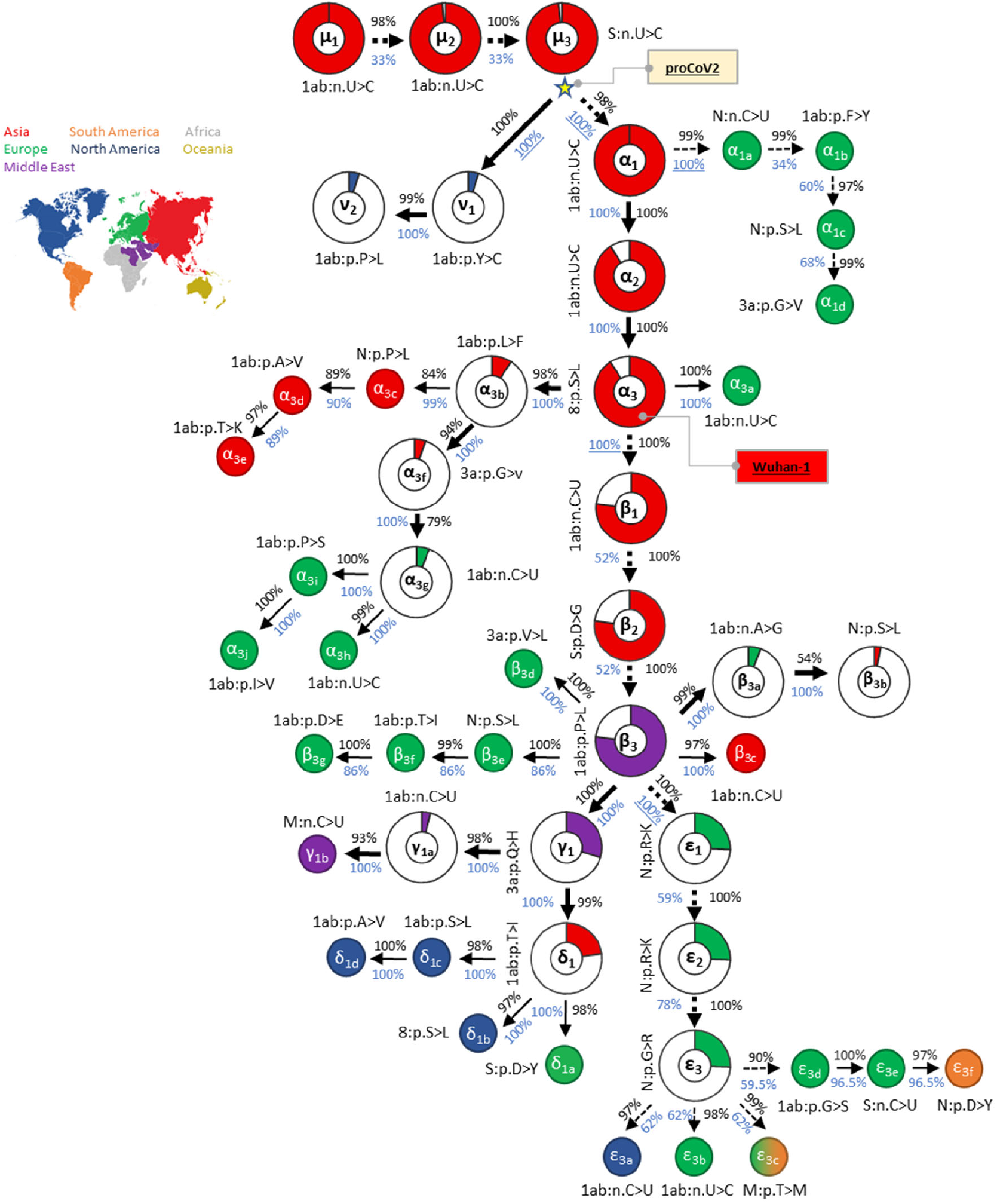
Mutational history graph of SARS-CoV-2 from the 29KG dataset. Thick arrows mark the pathway of widespread variants (frequency, *vf* ≥ 3%), and thin arrows show paths leading to other common mutations (3% > *vf* > 1%). The pie-charts’ size is proportional to variant frequency in the 29KG dataset, with pie-charts shown for variants with *vf* > 3% and pie color based on the world’s region where that mutation was first observed. A circle is used for all other variants, with the filled color corresponding to the earliest sampling region. The co-occurrence index (COI, black font) and the bootstrap confidence level (BCL, blue font) of each mutation and its predecessor mutation are shown next to the arrow connecting them. Underlined BCL values mark variant pairs for which BCLs were estimated for groups of variants (see *Methods*) because of the episodic nature of variant accumulation within groups resulting in lower BCLs (<80%; dashed arrows). Base changes (n.) are shown for synonymous mutations, and amino acid changes (p.) are shown for nonsynonymous mutations along with the gene/protein names (“ORF” is omitted from gene name abbreviations given in **Extended Data Table 1**). More details on each mutation are presented in **Extended Data Table 1**.

**Fig. 2.**
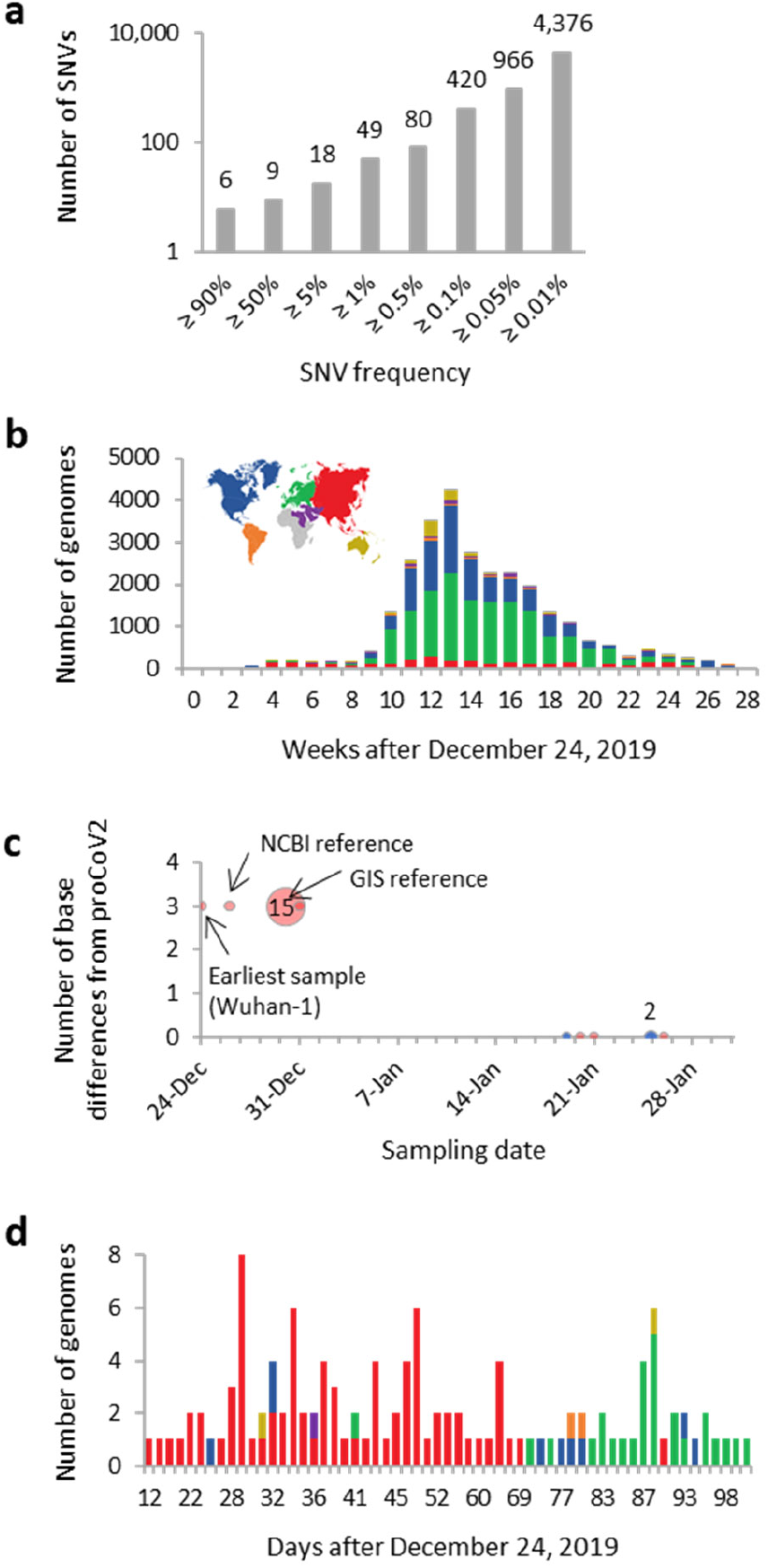
Counts of single nucleotide variants (SNVS) and genomes in the 29KG dataset. (**a**) Cumulative count of SNVs presented in the 29KG genome dataset at different frequencies. (**b**) The number of genomes in the 29KG collection that were isolated weekly during the pandemic. (**c**) The number of base differences from proCoV2 for genomes that were sampled in December 2019 and January 2020. The 18 genomes sampled in December 2019 in China (red) have three common SNVs different from proCoV2. In contrast, six genomes sampled in January 2020 in China (Asia, red) and the US (North America, blue) show no base differences. Multiple genomes (2 and 15) were sampled on two different days. (**d**) Temporal and spatial distribution of strains identical to proCoV2 at the protein sequence level, i.e., they have only μ mutations. The color scheme used to mark sampling locations is shown in panel b.

We found that new variants occurred in the genomic background of the variants preceding them in the reconstructed mutation history with a very high propensity (co-occurrence index, COI > 96.7%; **Fig. 1**). This suggests a strong signal to infer a sequential mutational history. Indeed, a bootstrap analysis involving genome resampling to assess the robustness of the mutation history produced high bootstrap confidence levels (BCLs) for key groups of mutations as well as many offshoots (**Fig. 1**; BCL > 95%). However, the order of some mutations was not established with a high BCL, e.g., the relative order of ε_1_, ε_2_, and ε_3_ mutations. This is because the three ε variants almost always occur together (7,624 genomes), and the intermediate combinations of ε variants occurred in only 42 genomes. Similarly, the count of genomes harboring all three β variants (22,739 genomes) far exceeded those with two or fewer β variants (201 genomes). There is a strong temporal tendency of variants to be sampled together (e.g., ε_1_ - ε_3_ and α_1a_-α_1d_), suggesting an episodic spread of variants (*P* << 0.01; see *Methods*). This episodic spreading of variants, which do not allow for determining the precise order of mutation appearance, may be caused by founder effects, positive selection, or both (e.g., ref.^28^). It may sometimes be an artifact of highly uneven regional and temporal genome sequencing that will produce a biased representative sample of the actual worldwide population (**Fig. 2b**).

### The progenitor genome

The root of the mutation tree is the most recent common ancestor (MRCA) of all the genomes analyzed, which gave rise to two early coronavirus lineages (ν and α; **Fig. 1**). The MRCA genome was the progenitor of all SARS-CoV-2 infections globally, henceforth proCoV2, and was likely carried by the first case of human transmission in the COVID-19 pandemic (index case)^20^. It existed on or before December 24, 2019, a date for which we have the sequence of SARS-CoV-2 infection in Wuhan, China (Wuhan-1; EPI_ISL 402123). A comparison of proCoV2 with Wuhan-1 genomes revealed three differences in the 49 positions, which was also true for other reference genomes (**Fig. 2c**). This suggests that the Wuhan-1 and the other earliest sampled genomes are derived coronavirus strains that arose from proCoV2 after the divergence of ν and α lineages (**Fig. 1**). The Wuhan-1 strain evolved by three successive α mutations in the progenitor (α_1_, α_2_, and α_3_), a progression that is statistically supported (BCL = 100%). This high resolution is made possible by 896 intermediate genomes containing one or two α variants in the 29KG dataset. Importantly, three closely-related non-human coronavirus genomes (bats and pangolin) all have the same base at these positions as does the proCoV2 genome, suggesting that the ancestral genome did not contain α variants. Furthermore, genomes with v variants of proCoV2 do not contain the other 47 variants, all of which occurred on the genomes containing α_1_-α_3_ that supports the inference that coronaviruses lacking α variants were the ancestors of Wuhan-1 and other genomes sampled in December 2019 in China (**Fig. 2c**). Therefore, we conclude that Wuhan-1 was not the direct ancestor of all the coronavirus infections globally.

Did proCoV2 propagate in the human population in 2020? A comparison of the proCoV2 genetic fingerprint (49 positions) in the 29KG collection revealed three matches in China (Fujian, Guangdong, and Hangzhou) and three in the US (Washington) in January 2020 (**Fig. 2c**). One more match was found in New York in March 2020, and the v mutant of proCoV2 was first sampled 59 days after the Wuhan-1 strain. This means that the progenitor coronavirus spread and mutated in the human population for weeks and months after the first reported COVID-19 cases.

Because proCoV2 is three bases different from the Wuhan-1 genome sampled on December 24, 2019, we estimate that the divergence of earliest variants of proCoV2 occurred 5.8 - 8.1 weeks prior based on the range of possible mutation rates of coronavirus genomes^20^. This timeline puts the presence of proCoV2 late-October to mid-November 2019 that is consistent with some other reports, including the report of a fragment of spike protein identical to Wuhan-1 in early December in Italy^18,20,29–31^. The sequenced segment of the spike protein is short (409 bases). It does not span positions in which 49 major early variants were observed, which means that the Italian Spike protein fragment can only confirm the existence of proCoV2 before the first coronavirus detection in China.

Comparisons of the protein sequences encoded by the proCoV2 genome revealed 131 other genomic matches, which contained only synonymous differences from proCoV2. A majority (89 genomes) of these matches were from coronaviruses sampled in China and other Asian countries (**Fig. 2d**). The first sequence was sampled 12 days after the earliest sampled virus, whose genome became available on December 24, 2019. Multiple matches were found in all sampled continents and detected as late as April 2020 in Europe. These spatiotemporal patterns suggest that proCoV2 already possessed the repertoire of protein sequences needed to infect, spread, and persist in the global human population (see also ref.^28^). Notably, none of these coronavirus genomes contained widely-studied Spike protein mutant (D614G), a β mutation that occurred in the genomes carrying all three α variants and was first seen in late January 2020.

We then analyzed a later snapshot of SARS-CoV-2 genome collection, consisting of genomes obtained from GISAID, acquired three months after the 29KG dataset. This dataset expanded the collection of coronavirus genomes from viral isolates collected after July 7, 2020 (16,739 genomes) and added 20,004 genome sequences from viral isolates dated before July 7, 2020. In the expanded MOA analysis, we retained 49 variants found with frequency > 1% in the 29KG dataset and added variants found with a frequency > 1% in the 68KG dataset (84 total variants; see **Extended Data Table 2**). MOA analysis of the 68KG dataset produced the proCoV2 genome identical to that inferred using the 29KG dataset (see *Methods*). We found one additional genome with a proCoV2 fingerprint sampled in Hubei, China, four weeks after the Wuhan-1 strain was reported.

The inferred mutation history from the 68KG dataset was well-supported with high COI and BCLs concordance with the mutation history produced using the 29KG dataset (**Fig. 3b**). Therefore, all the inferences reported for the 29KG dataset were robust to the expanded sampling of genomes. In the expanded mutation history, two new groups of variants were identified (ζ and η), which originated in mid-March 2020 and are found in relatively high frequency in the 68KG dataset (~4.4% and 8.0%, respectively; **Extended Data Table 2**). Variants in ζ and η groups also showed episodic accumulation of mutations, e.g., the count of genomes containing three ζ mutations (ζ_1_-ζ_3_; 2,955 genomes) was much larger than those with a subset of these variants (148 genomes). The episodic nature of mutational spread for 84 variants in the 68KG is statistically significant (*P* < 10^-8^), i.e., clusters of mutations together have become common variants (see *Methods*).

**Fig. 3.**
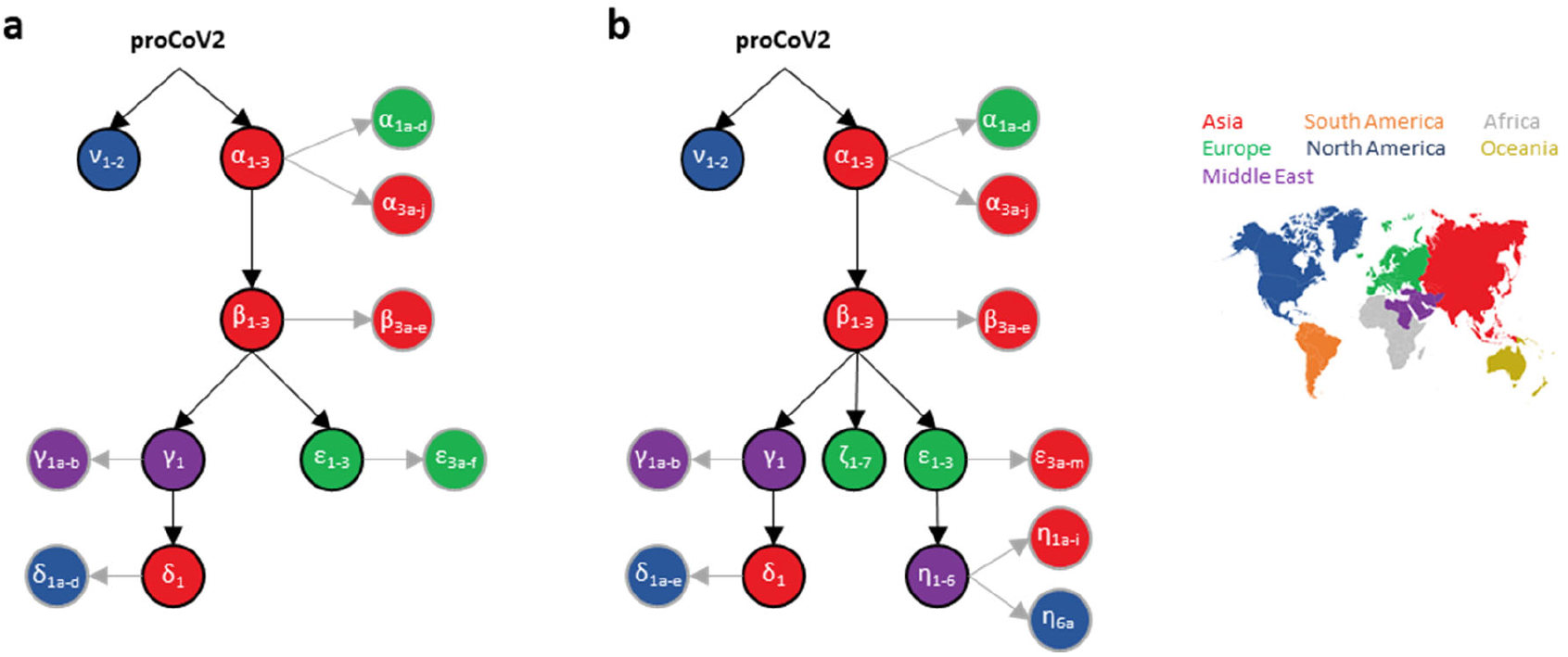
The backbone of SARS-CoV-2 mutational history. The mutational history inferred was from (**a**) 29KG and (**b**) 68KG datasets. Major variants and their mutational pathways are shown in black, and minor variants and their mutational pathways are gray. Circle color marks the region where variants were sampled first. The 68KG dataset contains 12 additional variants and more than two times the genomes than the 29KG dataset.

### Coronavirus fingerprints and spatiotemporal tracking

The progression of mutations in the mutation history directly transforms into a collection of genetic fingerprints or signatures. Each fingerprint represents a genome type containing all the variants on the path from that node up to the progenitor proCoV2. These fingerprints can classify genomes and track spatiotemporal patterns of dominant lineages genomes (see *Methods*). We use a shorthand to refer to each barcode in which only the major variant type is used. For example, **α** fingerprint refers to genomes that one or more of the α variants and no other major variants, and **αβ** fingerprint refers to genomes that contain at least one α, at least one β variant, and no other major variants. This nomenclature is intuitive and provides a way to glean evolutionary information from the coronavirus lineage’s name. In the 68KG dataset (October 12, 2020 GISAID snapshot), global frequencies of major proCoV2 fingerprints were **αβε** (32.1%), **αβγδ** (17.7%), **αβ** (16.7%), **αβεη** (9.9%), **αβ (**9.8%), **αβγ** (6.8%), **αβζ** (4.5%), and **ν** (2.4%).

**Figure 4** shows the evolving spatiotemporal of all major fingerprints in Asia, Europe, and North America inferred for an expanded dataset of 172,480 genomes (December 30, 2020 snapshot). Spatiotemporal patterns in cities, countries, and other regions are available online at http://sars2evo.datamonkey.org/. We observe the spread and replacement of prevailing strains in Europe (**αβε** with **αβζ**) and Asia (**α** with **αβε**), the preponderance of the same strain for most of the pandemic in North America (**αβγδ**), and the continued presence of multiple high-frequency strains in Asia and North America. Spatiotemporal patterns of strain spread converged for Europe and Asia by July-August 2020 to **αβε** genetic fingerprints. These patterns diverged from North America, where **αβ** along with its mutant (**αβγδ**) were common. After that, Europe saw **ζ** variants of **αβ** grow (**αβζ**), replacing **αβε** genomes and its new **η** offshoot (**αβεη**) (e.g., ref.^32^). The **ζ** mutations were first detected three weeks after the sampling of the first **ε** variants. Remarkably, **αβγδ** has remained the dominant lineage in North America since April 2020, in contrast to the turn-over seen in Europe and Asia. More recently, novel fast-spreading variants have been reported (e.g., ref^33^). In particular, an S protein variant (N501Y) from South Africa and London has rapidly increased^33^. Coronaviruses with N501Y variant in South Africa carry the **αβγδ** genetic fingerprint, whereas those in London carry the **αβε** genetic fingerprint. This means that the N501Y mutation arose independently in two coronavirus lineages that show convergent patterns of increased spread. At present, **αβζ** dominates the UK, and the number of genomes publicly available from South Africa is relatively small to make reliable inferences at present (see http://sars2evo.datamonkey.org for future updates). Overall, our mutational fingerprinting and nomenclature provides a simple way to glean the ancestry of new variants in contrast to phylogenetic designations (e.g., B.1.350 and B.1.1.7^33^).

**Fig. 4.**
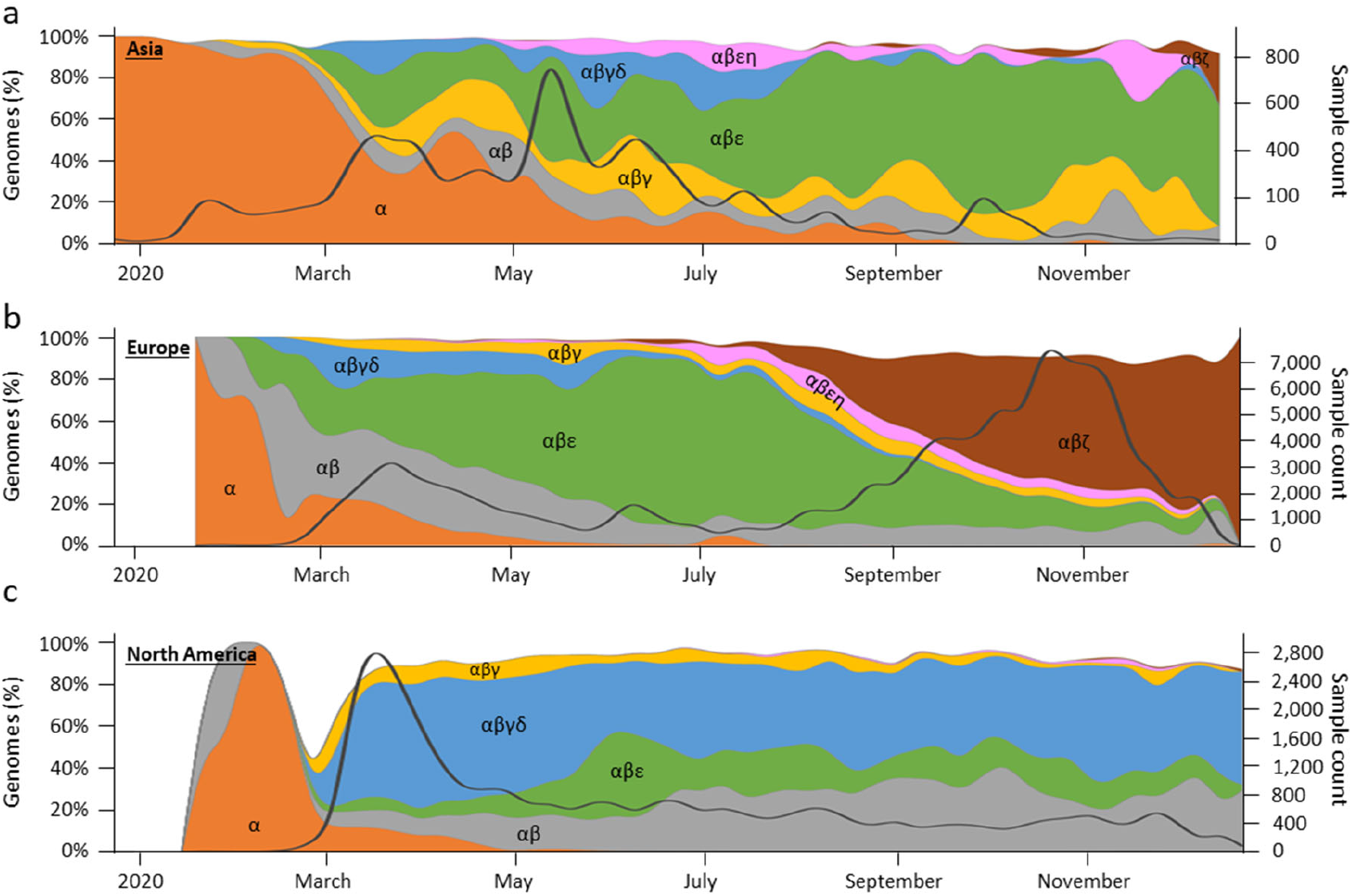
Spatiotemporal dynamics of 172,480 SARS-CoV-2 genomes (December 2019-2020). Spatiotemporal patterns of genomes mapped to lineages containing different combinations of major variants in (**a**) Asia, (**b**) Europe, and (**c**) North America. The number of genomes mapped to major variant lineages contains all of its offshoots, e.g., α lineage contains all the genomes with α_1_ – α_3_, α_1a_ – α_1d_, and α_3a_ – α_3j_ variants only. The stacked graph area is the proportion of genomes mapped to the corresponding lineage. The solid black line shows the count of total genome samples. Spatiotemporal patterns in cities, countries, and other regions are available online at http://sars2evo.datamonkey.org/.

## Conclusions

Through innovative analyses of two large collections of SARS-CoV-2 genomes, we have consistently reconstructed the same progenitor coronavirus genome and identified its presence worldwide for many months after the pandemic began. The progenitor genome is a better reference for rooting phylogenies, orienting mutations, and estimating sequence divergences. The reconstructed mutational history of SARS-CoV-2 revealed major mutational fingerprints to identify and track the novel coronavirus’s spatiotemporal evolution, revealing convergences and divergences of dominant strains among geographical regions from an analysis of more than 174 thousand genomes.

Furthermore, the approach taken here to reconstruct the progenitor genome and discover key mutational events will generally be applicable for analyzing pathogens during the early stages of outbreaks. The approach is scalable for even bigger datasets because it does not require more phylogenetically informative variants with an increasing number of samples. In fact, it benefits from bigger datasets as they afford more accurate estimates of individual and co-occurrence frequencies of variants and enable more reliable detection of lower frequency variants. Its continued application to SARS-CoV-2 genomes and other pathogen outbreaks will produce their ancestral genomes and their spatiotemporal dynamics, improving our understanding of the past, current, and future evolution of pathogens and associated diseases.

## Methods

### Genome data acquisition and processing

We first downloaded 60,332 SARS-CoV-2 genomes from the GISAID^27^ database, along with information on sample collection dates and locations (until July 7, 2020). Of all the genomes downloaded, we only retained those with greater than 28,000 bases and were marked as originating from human hosts and passing controls detailed below. Similarly, the second dataset, the 68KG dataset, was assembled from 133,741 genomes and downloaded on October 12, 2020. Again, we retained only those with greater than 28,000 bases and marked as originating from human hosts.

Each genome was subjected to codon-aware alignment with the NCBI reference genome (accession number NC_045512) and then subdivided into ten regions based on CDS features: ORF1a (including nsp10), ORF1b (starting with nsp12), S, ORF3a, E, M, ORF6, ORF7a, ORF8, N, and ORF10. Gene ORF7b was removed because it was too short for alignment and comparisons. For each region, we scanned and discarded sequences containing too many ambiguous nucleotides to remove data with too many sequencing errors. Thresholds were 0.5% for the S gene, 0.1% for ORF1a and ORF1b genes, and 1% for all other genes. We mapped individual sequences to the NCBI reference genome (NC_045512) using a codon-aware extension to the Smith-Waterman algorithm implemented in HyPhy^34^ (https://github.com/veg/hyphy-analyses/tree/master/codon-msa), translated mapped sequence to amino-acids, and performed multiple protein sequence alignment with the auto settings function of MAFFT (version 7.453)^35^. Codon sequences were next mapped onto the amino-acid alignment. The multiple sequence alignment of SARS-CoV-2 genomes was aligned with the sequence of three closest outgroups, including the coronavirus genomes of the *Rhinolophus affinis* bat (RaTG13), *R. malayanus* bat (RmYN02), and *Manis javanica* pangolin (MT121216.1)^36,37^. The alignment was visually inspected and adjusted in Geneious Prime 2020.2.2 (https://www.geneious.com). The final alignment contained all genomic regions except ORF7b and non-coding regions (5’ and 3’ UTRs, and intergenic spacers). After these filtering and alignment steps, the multiple sequence alignment contained 29,115 sites and 29,681 SARS-CoV-2 genomes for the July 7, 2020 snapshot, which we refer to as the 29KG dataset. For the October 12 snapshot, there were 68,057 sequences, which we refer to as the 68KG dataset. We also conducted a spatiotemporal analysis on an expanded dataset containing 172,480 genomes (172KG) acquired on December 30, 2020.

### Reference genomes and collection dates

We used the dates of viral collections provided by the GISAID database^27^ in all our analyses if they were resolved to the day (i.e., we discarded data that only contained partial dates, e.g., April 2020). All genomes were used in the mutation ordering analyses, but genomes with incomplete sampling dates were excluded from the spatiotemporal analyses and derived interpretations. We noted that the earliest sample included in GISAID (ID: EPI_ISL_402123) was collected on December 24, 2019, although the NCBI website lists its collection date as December 23, 2019 (GenBank ID: MT019529). Therefore, we used the GISAID collection date for the sake of consistency. Regarding the NCBI reference genome (GenBank ID: NC_045512; GISAID ID: EPI_ISL_402125)^38^, this sample was collected on December 26, 2019^39^. We also used the GIS reference genome in our analysis (ID: EPI_ISL_402124), collected on December 30, 2019^40^.

### Mutation order analyses (MOA)

First, we analyzed the 29KG dataset. We used a maximum likelihood method, SCITE^23^, and variant co-occurrence analyses for reconstructing the order of mutations corresponding to 49 common variants (frequency > 1%) observed in this dataset. MOA has demonstrated high accuracy for analyzing tumor cell genomes that reproduce clonally, have frequent sequencing errors, and exhibit limited sequence divergence^23,24^. In MOA, higher frequency variants are expected to have arisen earlier than low-frequency variants in clonally reproducing populations^23,26^. We used the highest frequency variants to anchor the analysis and the shared co-occurrence of variants among genomes to order mutations while allowing probabilistically for sequencing errors and pooled sequencing of genomes^23^. MOA is different from traditional phylogenetic approaches where positions are treated independently, i.e., the shared co-occurrence of variants is not directly utilized in the inference procedure. Notably, both traditional phylogenetic and mutation order analyses are expected to produce concordant patterns when sequencing errors and other artifacts are minimized. However, sequencing errors and limited mutational input during the coronavirus history adversely impact traditional methods, as does the fact that the closest coronaviruses useable as outgroups have more than a thousand base differences from SARS-CoV-2 genomes that only differ in a handful of bases from each other^7,9,10^.

MOA requires a binary matrix of presence/absence (1/0) of mutants, which is straightforward in analyzing cell sequences from tumors because they arise from normal cells that supply the definitive ancestral state. To designate mutation orientations for applying MOA in SARS-CoV-2 analysis, we devised a simple approach in which we began by comparing nucleotides at the 49 genomic positions among three closely-related genomes (bat RaTG13, bat RmYN02, and pangolin MT121216.1)^41^. We chose the consensus base to be the initial reference base, such that SARS-CoV-2 genome bases were coded to be “0” whenever they were the same as the consensus base at their respective positions. All other bases were assigned a “1.” There were 39 positions in which all three outgroup genomes were identical to each other and 9 in which two of the outgroups showed the same base. In the remaining position (28657), all three outgroups differed, so we selected the base found in the gene with the highest sequence similarity to the human SARS-CoV-2 NCBI reference genome (NC_045512) because SARS-CoV-2’s ancestor likely experienced genomic recombination before its zoonotic transfer into humans^28,42,43^. At one position, both major and minor bases in humans were different from the consensus base in the outgroups, so we assigned the mutant status to the minority base (U; *vf* = 29.8%). All missing and ambiguous bases were coded to be ignored (missing data) in all the analyses.

These initially assigned mutation orientations were tested in a subsequent investigation of variants’ cooccurrence index (COI). COI for a given variant (*y*) is the number of genomes that contain *y* and its directly preceding mutation (*x*) in the mutation history, divided by the number of genomes that contain *y*. When COI was lower than 70%, we reversed each position’s mutation orientation individually and selected the mutation orientation that produced mutation histories with the highest COI.

In the SCITE analysis of 49 variants and 29,861 genomes, we started with default parameter settings of false-negative rate (FNR = 0.21545) and false-positive rate (FPR = 0.0000604) of mutation detection. We carried out five independent runs to ensure stability and convergence to obtain 29KG collection-specific estimates of FNR and FPR by comparing the observed and predicted sequences based on this mutation graph. The estimated FNR (0.00488) and FPR (0.00800) were very different from the SCITE default parameters, where the estimated FNR was much lower. This difference in error rates is unsurprising because we used only common variants (*vf* > 1%), and the 29KG dataset was not obtained from single-cell sequencing in which dropout during single-cell tumor sequencing elevates FNR, i.e., mutant alleles are not sequenced.

As noted above, the initial mutation orientations were simply the starting designations for our analysis, which are subsequently investigated by evaluating the COI of each variant in the reconstructed mutation history. In this process, we reverse ancestor/mutant coding for variants that received low COI to examine if a mutation history with higher COI can be generated. Two positions (3037 and 28854) received low COI (<70%). At position 3037, the reversed encoding (C→U) received significantly higher COI (100%) than the starting encoding (U→C; 60%), so the position was recoded. At position 28854, the ordering and direction of mutation remained ambiguous despite extensive analyses, but it did not impact the predicted MRCA sequence. Therefore, we only recoded the column for position 3037 and generated a new 49 × 29861 (SNVs × genomes) matrix to conduct a SCITE analysis.

At one position (28657), all three outgroup sequences had different bases, so we initially selected the base found in the gene with the highest sequence similarity to the human SARS-CoV-2 NCBI reference genome. We next tested if reversed encoding produced a better mutation graph. The reversed encoding produced a mutation graph with a much higher log-likelihood (−32355.58 and −30289.92, for the initial and reversed encoding, respectively; *P* << 0.01 using the AIC protocol in ref.^44^). Therefore, we recoded position 28657 and generated a new 49 × 29861 (SNVs × genomes) matrix.

It was subjected to SCITE analysis and produced a mutation graph for 49 variants in the 29KG dataset. This graph predicts an FNR of 0.00418 and FPR of 0.00295 per base. Using these new FNR and FPR, we again performed SCITE analysis and produced the final mutation history graph. Starting from the top of a mutation graph, a distinct Greek symbol was assigned to a group of mutations that were occurred sequentially, and variants with similar frequency were assigned the same Greek symbol (μ, ν, α, β, γ, δ, and ε). The high-frequency variants with the same Greek symbol were distinguished by numbers to represent the sequential relationship, e.g., α_1_ and α_2_. When an offshoot of a high-frequency mutation had low variant frequency, we assigned it the same Greek symbol and number to represent the parent-offspring relationship and further distinguished descendants by adding a small letter, e.g., α_1a_ and α_1b_.

In this mutation graph, the most recent common ancestor (MRCA) corresponds to the progenitor that gave rise to ν and α lineages. MRCA is the progenitor of all human SARS-CoV-2 infections (proCoV2), which descended from the parental lineage that divergence form and its closest relatives, including bats and pangolins. We estimate that proCoV2 existed 5.8 to 8.1 weeks before December 24, 2019, on which the Wuhan-1 was sampled, by using SARS-CoV-2 HPD mutation rate range of 6.64×10^-4^ – 9.27×10^-4^ substitutions per site per year^20^. We have made available the proCoV2 genome sequence in FastA format at http://igem.temple.edu/COVID-19, which is the same as the NCBI reference genome with base differences corresponding to α_1_ – α_3_ mutations at positions 18060, 8782, and 28144, as discussed in the main text. In this mutation graph, COI for each variant is shown next to the arrow.

### Bootstrap analysis

We assessed the robustness of the mutation history inference to genome sampling by bootstrap analysis. We generated 100 bootstrap replicate datasets, each built by randomly selecting 29,861 genomes with replacement. Then, SCITE was used to infer the mutation graph for each replicate dataset. Bootstrap confidence level, scored for each variant pair, was the number of replicates in which the given pair of variants were directly connected in the mutation history in the same way as shown in **figure 1**. BCLs were often lower for major variants within groups (e.g., ε_1_ – ε_3_) because they occur with very similar frequencies. This feature adversely affected the BCL values of mutation orders between groups, e.g., β and ε. In this case, we considered each group as a single entity. We computed BCL to be the proportion of replicates in which pairs of groups were directly connected in the mutation history in the same way as shown in **figure 1**. Groups used were β_1_-β_3_, ε_1_-ε_3_, and α_1a_-α_1d_. All of these BCL values are shown with an underline.

### Temporal concordance

Because mutation ordering analysis analyses did not use spatial or temporal information for genomes or mutations, it can be validated by evaluating the concordance of the inferred order of mutations with the timing of their first appearance (*tf*). Using the genomes for which virus sampling day, month, and year were available, we determined *tf* for every variant in the 29KG dataset. For a mutation *i*, we compared its *tf*(*i*) with *tf*(*j*) such that *j* is the nearest preceding mutation in the mutation graph. We found that *tf*(*j*) ≥ *tf*(*i*) for 47 of 49 mutations, except for β_3b_ and β_3c_ pairs. These two offshoot mutants of β_3_ were sampled 35 days (β_3b_) and 12 days (β_3c_) earlier than their predecessors, which could be due to their low frequency or sequencing error. COI of one variant (β_3b_) was low (54%), but the other variant (β_3c_) had a very high COI (97%).

### Mutational fingerprints

Each node in the mutational history graph predicts an intermediate (ancestral) or a tip sequence, containing all the mutations from that node to the mutation graph’s root. The mutational fingerprint is then produced directly from the mutation history graph drawn as a directional graph anchored on the root node. We compared our mutational fingerprints of the genomes in the 29KG dataset with a phylogeny-based classification^1^ obtained using the Pangolin service (v2.0.3; https://pangolin.cog-uk.io/). We assigned each of the 29K genomes to a fingerprint based on the highest sequence similarity at positions containing 49 common variants. Mismatches were allowed, as sequencing errors could create them. A small fraction of genomes (1.8%) could not be assigned unambiguously to one fingerprint, so they were excluded and investigated in the future. The number of genomes assigned to each fingerprint is shown in **Extended Data Table 1**. We submitted genome sequences to the Pangolin website for classification one-by-one, and a clade designation was received. The results are summarized in **Extended Data Figure 1**. In this table, all phylogenetic-groups with fewer than 20 genomes were excluded.

Of the 80 phylogenetic groups shown, 74 are defined primarily by a single mutation-based fingerprint, as more than 90% of the genomes in those phylogenetic groups share the same fingerprint. This includes all small and medium-sized phylogenetic groups (up to 488 genomes) and two large groups (A.1 with 1,377 genomes and B.1.2 with 749 genomes). One large group, B.1.1, predominately connects with ε_3_ node (79%, 4,832 genomes), but some of its members belong to ε_3_ offshoots because they contain respective diagnostic mutations. For group B.1.1.1, two other ε_3_ offshoots are mixed up almost equally. Three other large differences between mutational fingerprint-based classification and phylogeny-based grouping are seen for A, B, B1.1, and B.2 groups. These differences are likely because the location of the root and the earliest branching order of coronavirus lineages are problematic in phylogeny-based classifications^7,9,10,14^. Overall, our mutational fingerprints are immediately informative about the mutational ancestry of genomes.

### Analysis of 68KG dataset

We repeated the above MOA procedure on the 68KG dataset (68,057 genomes). This 68KG data contained 72 common variants (>1% frequency). For direct comparison purposes, we added 12 variants that were common variants on 29KG data, but their frequency had become less than 1% in the 68KG data. Therefore, we used 84 variants in total and constructed a matrix of 84 × 68,057 (SNVs × genomes) for the SCITE analysis to determine the mutational order. We also conducted the bootstrap analysis and assigned mutational fingerprints using the procedure mentioned above. The number of genomes mapped to each fingerprint is listed in **Extended Data Table 2**.

### Spatiotemporal analysis of 172KG dataset

We developed a sequence classification protocol that first calls variants in a viral genome using proCoV2 as the reference sequence using minimap2^45^. Then, it assigns the sequence to a path in the mutation graph with the highest concordance (Jaccard index). It is implemented in a simple browser-based tool, which shows the example output for ENA accession number MT675945 (**Extended Data Figure 2**; http://sars2evo.datamonkey.org). The classification is conducted on the client-side such that the researcher’s data never leaves their personal computer.

### Testing episodic spread of variants

We performed non-parametric Wald–Wolfowitz runs-tests^46,47^ of the null hypothesis that the first sampling of variants is randomly distributed over time (i.e., evenly spaced). The null hypothesis was rejected for both 29KG and 64KG analysis at *P* << 0.01, suggesting significant temporally clustering in both 29KG dataset and 64KG datasets. Because many mutations were first sampled on December 24, 2019, we only included one mutation for that day to avoid biasing the test.

### Data Availability and Code Availability

Live evolutionary history and spatiotemporal distributions of common variants can be accessed via http://igem.temple.edu/COVID-19 (beta version). All genome sequences and metadata are available publicly at GISAID (https://www.gisaid.org/), and the predicted proCoV2 sequence is available at http://igem.temple.edu/COVID-19. The other relevant information is provided in the supplementary materials.

## Acknowledgments

We thank all the authors and organizations who have kindly deposited and shared genome data on GISAID (see http://igem.temple.edu/COVID-19 for a list of all the authors). We thank Ananias Escalante, Rob Kulathinal, Li Liu, Jose Barba-Montoya, Antonia Chroni, Ravi Patel, and Caryn Babaian for their critical comments. We appreciate the technical support provided by Jared Knoblauch and Glen Stecher. This research was supported by grants from the U.S. National Science Foundation to S.K. (GCR-1934848, DEB-2034228) and S.P. (DBI-2027196) and from the U.S. National Institutes of Health to S.K. (GM-0126567-03 and 139504-01) and S.P. (AI-134384).

## Author Contributions

S.K. and S.M. conceived the project, designed analyses and visualizations, conducted initial analyses, and wrote the manuscript. S.P., S.W., and S.K. designed and developed the browser resource and tools. S.P., S.W., and M.A.C.O. assembled sequence alignments. M.A.C.O., S.M., S.S., and Q.T. conducted analyses and rendered visualizations. All authors intellectually contributed by discussing results and patterns, and everyone contributed to writing the manuscript.

## Additional Information

**Supplementary Information** is available for this paper. Correspondence and requests for materials should be addressed to.

---

**Extended Data Table 1.** SARS-CoV-2 variants in 29KG dataset.

| Mutant (major) | Mutant (minor) | Gene | Genomic Position | Nucleotide change | Amino acid change | Time (days) | Variant Frequency | Genomes mapped | First location |
| --- | --- | --- | --- | --- | --- | --- | --- | --- | --- |
| μ<sub>1</sub> |  | ORF1ab | 2416 | U>C |  | 0 | 98.1% | 0 | China, Asia |
| μ<sub>2</sub> |  | ORF1ab | 19524 | U>C |  | 0 | 98.6% | 0 | China, Asia |
| μ<sub>3</sub> |  | S | 23929 | U>C |  | 0 | 98.4% | 18 | China, Asia |
| α<sub>1</sub> |  | ORF1ab | 18060 | U>C |  | 0 | 95.1% | 849 | China, Asia |
|  | α<sub>1a</sub> | N | 28657 | C>U |  | 63 | 1.3% | 2 | France, Europe |
|  | α<sub>1b</sub> | ORF1ab | 9477 | U>A | F>Y | 63 | 1.2% | 3 | France, Europe |
|  | α<sub>1c</sub> | N | 28863 | C>U | S>L | 63 | 1.2% | 5 | France, Europe |
|  | α<sub>1d</sub> | ORF3a | 25979 | G>U | G>V | 63 | 1.2% | 344 | France, Europe |
| α<sub>2</sub> |  | ORF1ab | 8782 | U>C |  | 0 | 91.0% | 47 | China, Asia |
| α<sub>3</sub> |  | ORF8 | 28144 | C>U | S>L | 0 | 90.8% | 1115 | China, Asia |
|  | α<sub>3a</sub> | ORF1ab | 1606 | U>C |  | 43 | 1.7% | 501 | United Kingdom, Europe |
|  | α<sub>3b</sub> | ORF1ab | 11083 | G>U | L>F | 24 | 9.2% | 376 | China, Asia |
|  | α<sub>3c</sub> | N | 28311 | C>U | P>L | 64 | 1.9% | 3 | South Korea, Asia |
|  | α<sub>3d</sub> | ORF1ab | 13730 | C>U | A>V | 71 | 1.8% | 3 | Taiwan/Malaysia, Asia |
|  | α<sub>3e</sub> | ORF1ab | 6312 | C>A | T>K | 71 | 1.7% | 483 | Taiwan/Malaysia, Asia |
|  | α<sub>3f</sub> | ORF3a | 26144 | G>U | G>V | 28 | 5.1% | 121 | China, Asia |
|  | α<sub>3g</sub> | ORF1ab | 14805 | C>U |  | 54 | 6.0% | 334 | United Kingdom, Europe |
|  | α<sub>3h</sub> | ORF1ab | 17247 | U>C |  | 64 | 2.0% | 580 | Switzerland, Europe |
|  | α<sub>3i</sub> | ORF1ab | 2558 | C>U | P>S | 54 | 1.7% | 26 | United Kingdom, Europe |
|  | α<sub>3j</sub> | ORF1ab | 2480 | A>G | I>V | 54 | 1.6% | 462 | United Kingdom, Europe |
| β<sub>1</sub> |  | ORF1ab | 3037 | C>U |  | 31 | 77.0% | 11 | China, Asia |
| β<sub>2</sub> |  | S | 23403 | A>G | D>G | 31 | 77.1% | 36 | China, Asia |
| β<sub>3</sub> |  | ORF1ab | 14408 | C>U | P>L | 41 | 76.9% | 3032 | Saudi Arabia, Middle East |
|  | β<sub>3a</sub> | ORF1ab | 20268 | A>G |  | 64 | 5.7% | 1213 | Italy, Europe |
|  | β<sub>3b</sub> | N | 28854 | C>U | S>L | 29 | 3.1% | 527 | China, Asia |
|  | β<sub>3c</sub> | ORF1ab | 15324 | C>U |  | 29 | 2.3% | 678 | China, Asia |
|  | β<sub>3d</sub> | ORF3a | 25429 | G>U | V>L | 77 | 1.7% | 485 | United Kingdom, Europe |
|  | β<sub>3e</sub> | N | 28836 | C>U | S>L | 74 | 1.6% | 3 | Switzerland, Europe |
|  | β<sub>3f</sub> | ORF1ab | 13862 | C>U | T>I | 74 | 1.6% | 50 | Switzerland, Europe |
|  | β<sub>3g</sub> | ORF1ab | 10798 | C>A | D>E | 86 | 1.4% | 414 | United Kingdom, Europe |
| γ<sub>1</sub> |  | ORF3a | 25563 | G>U | Q>H | 41 | 29.8% | 884 | Saudi Arabia, Middle East |
|  | γ<sub>1a</sub> | ORF1ab | 18877 | C>U |  | 41 | 4.0% | 757 | Saudi Arabia, Middle East |
|  | γ<sub>1b</sub> | M | 26735 | C>U |  | 41 | 1.5% | 439 | Saudi Arabia, Middle East |
| δ<sub>1</sub> |  | ORF1ab | 1059 | C>U | T>I | 54 | 23.0% | 5157 | Singapore, Asia |
|  | δ<sub>1a</sub> | S | 24368 | G>U | D>Y | 75 | 1.3% | 389 | Sweden, Europe |
|  | δ<sub>1b</sub> | ORF8 | 27964 | C>U | S>L | 76 | 2.7% | 790 | USA, North America |
|  | δ<sub>1c</sub> | ORF1ab | 11916 | C>U | S>L | 72 | 1.6% | 166 | USA, North America |
|  | δ<sub>1d</sub> | ORF1ab | 18998 | C>U | A>V | 72 | 1.0% | 305 | USA, North America |
| ε<sub>1</sub> |  | N | 28881 | G>A | R>K | 54 | 25.7% | 2 | United Kingdom, Europe |
| ε<sub>2</sub> |  | N | 28882 | G>A | R>K | 54 | 25.7% | 2 | United Kingdom, Europe |
| ε<sub>3</sub> |  | N | 28883 | G>C | G>R | 54 | 25.7% | 5365 | United Kingdom, Europe |
|  | ε<sub>3a</sub> | ORF1ab | 313 | C>U |  | 66 | 2.1% | 608 | USA, North America |
|  | ε<sub>3b</sub> | ORF1ab | 19839 | U>C |  | 64 | 1.5% | 452 | Switzerland, Europe |
|  | ε<sub>3c</sub> | M | 27046 | C>U | T>M | 69 | 1.6% | 453 | Worldwide |
|  | ε<sub>3d</sub> | ORF1ab | 10097 | G>A | G>S | 69 | 2.5% | 5 | Denmark, Europe |
|  | ε<sub>3e</sub> | S | 23731 | C>U |  | 69 | 2.5% | 403 | Denmark, Europe |
|  | ε<sub>3f</sub> | N | 28580 | G>U | D>Y | 69 | 1.2% | 353 | Chile, South America |
| ν<sub>1</sub> |  | ORF1ab | 17858 | A>G | Y>C | 59 | 4.7% | 32 | USA, North America |
| ν<sub>2</sub> |  | ORF1ab | 17747 | C>U | P>L | 59 | 4.7% | 1374 | USA, North America |

Note.- Genomic locations correspond to those of the NCBI genome (GenBank ID: NC\_04551.2). Amino acid changes are shown for nonsynonymous variants.

**Extended Data Table 2.** SARS-CoV-2 variants in the 68KG dataset.

| Mutant (major) | Mutant (minor) | Gene | Genomic Position | Nucleotide change | Amino acid change | Time (days) | Variant Frequency | Genomes mapped | First location |
| --- | --- | --- | --- | --- | --- | --- | --- | --- | --- |
| $\mu_1$ | | ORF1ab | 2416 | U>C | | 0 | 98.4% | 0 | China, Asia |
| $\mu_2$ | | ORF1ab | 19524 | U>C | | 0 | 99.0% | 18 | China, Asia |
| $\mu_3$ | | S | 23929 | U>C | | 0 | 98.9% | 0 | China, Asia |
| $\mu_4$ | | ORF1ab | 15933 | U>C | | 0 | 98.8% | 0 | China, Asia |
| $\mu_5$ | | ORF8 | 27944 | U>C | | 0 | 97.0% | 0 | China, Asia |
| $\mu_6$ | | ORF1ab | 6286 | U>C | | 0 | 95.6% | 0 | China, Asia |
| $\mu_7$ | | S | 22444 | U>C | | 0 | 98.7% | 0 | China, Asia |
| $\alpha_1$ | | ORF1ab | 18060 | U>C | | 0 | 97.3% | 1114 | China, Asia |
| | $\alpha_{1a}$ | N | 28657 | C>U | | 63 | 1.0% | 3 | France, Europe |
| | $\alpha_{1b}$ | ORF1ab | 9477 | U>A | F>Y | 63 | 0.7% | 3 | France, Europe |
| | $\alpha_{1c}$ | N | 28863 | C>U | S>L | 63 | 0.7% | 7 | France, Europe |
| | $\alpha_{1d}$ | ORF3a | 25979 | G>U | G>V | 63 | 0.7% | 451 | France, Europe |
| $\alpha_2$ | | ORF1ab | 8782 | U>C | | 0 | 94.9% | 51 | China, Asia |
| $\alpha_3$ | | ORF8 | 28144 | C>U | S>L | 0 | 94.9% | 1281 | China, Asia |
| | $\alpha_{3a}$ | ORF1ab | 1606 | U>C | | 43 | 0.9% | 578 | United Kingdom, Europe |
| | $\alpha_{3b}$ | ORF1ab | 11083 | G>U | L>F | 24 | 7.5% | 417 | China, Asia |
| | $\alpha_{3c}$ | N | 28311 | C>U | P>L | 64 | 1.4% | 4 | South Korea, Asia |
| | $\alpha_{3d}$ | ORF1ab | 13730 | C>U | A>V | 33 | 1.4% | 5 | China, Asia |
| | $\alpha_{3e}$ | ORF1ab | 6312 | C>A | T>K | 71 | 1.2% | 767 | Taiwan, Asia |
| | $\alpha_{3f}$ | ORF3a | 26144 | G>U | G>V | 28 | 3.0% | 160 | China, Asia |
| | $\alpha_{3g}$ | ORF1ab | 14805 | C>U | | 54 | 3.7% | 511 | United Kingdom, Europe |
| | $\alpha_{3h}$ | ORF1ab | 17247 | U>C | | 64 | 1.0% | 682 | Switzerland, Europe |
| | $\alpha_{3i}$ | ORF1ab | 2558 | C>U | P>S | 54 | 1.0% | 44 | United Kingdom, Europe |
| | $\alpha_{3j}$ | ORF1ab | 2480 | A>G | I>V | 54 | 1.0% | 648 | United Kingdom, Europe |
| $\beta_1$ | | ORF1ab | 3037 | C>U | | 31 | 87.2% | 45 | China, Asia |
| $\beta_2$ | | S | 23403 | A>G | D>G | 31 | 87.2% | 15 | China, Asia |
| $\beta_3$ | | ORF1ab | 14408 | C>U | P>L | 41 | 87.1% | 4450 | Saudi Arabia, Middle East |
| | $\beta_{3a}$ | ORF1ab | 20268 | A>G | | 64 | 6.0% | 2388 | Italy, Europe |
| | $\beta_{3b}$ | N | 28854 | C>U | S>L | 29 | 4.5% | 1782 | China, Asia |
| | $\beta_{3c}$ | ORF1ab | 15324 | C>U | | 29 | 2.2% | 1463 | China, Asia |
| | $\beta_{3d}$ | ORF3a | 25429 | G>U | V>L | 77 | 1.1% | 719 | United Kingdom, Europe |
| | $\beta_{3e}$ | N | 28836 | C>U | S>L | 74 | 0.8% | 3 | Switzerland, Europe |
| | $\beta_{3f}$ | ORF1ab | 13862 | C>U | T>I | 74 | 0.8% | 85 | Switzerland, Europe |
| | $\beta_{3g}$ | ORF1ab | 10798 | C>A | | 86 | 0.6% | 435 | United Kingdom, Europe |
| $\gamma_1$ | | ORF3a | 25563 | G>U | Q>H | 41 | 24.4% | 1671 | Saudi Arabia, Middle East |
| | $\gamma_{1a}$ | ORF1ab | 18877 | C>U | | 41 | 4.2% | 1201 | Saudi Arabia, Middle East |
| | $\gamma_{1b}$ | M | 26735 | C>U | | 41 | 2.7% | 1784 | Saudi Arabia, Middle East |
| $\delta_1$ | | ORF1ab | 1059 | C>U | T>I | 54 | 17.6% | 8284 | Singapore, Asia |
| | $\delta_{1a}$ | S | 24368 | G>U | D>Y | 75 | 0.7% | 466 | Sweden, Europe |
| | $\delta_{1b}$ | ORF8 | 27964 | C>U | S>L | 76 | 2.9% | 1152 | USA, North America |
| | $\delta_{1c}$ | ORF1ab | 11916 | C>U | S>L | 72 | 1.9% | 807 | USA, North America |
| | $\delta_{1d}$ | ORF1ab | 18998 | C>U | A>V | 72 | 0.7% | 458 | USA, North America |
| | $\delta_{1e}$ | ORF1ab | 10319 | C>U | L>F | 76 | 1.2% | 799 | USA, North America |
| $\zeta_1$ | | ORF1ab | 445 | U>C | | 179 | 4.4% | 18 | Netherlands, Europe |
| $\zeta_2$ | | M | 26801 | C>G | | 82 | 4.3% | 7 | Canada, North America |
| $\zeta_3$ | | S | 22227 | C>U | A>V | 84 | 4.5% | 1 | Spain, Europe |
| $\zeta_4$ | | N | 28932 | C>U | A>V | 96 | 4.4% | 5 | Portugal, Europe |
| $\zeta_5$ | | ORF10 | 29645 | G>U | V>L | 78 | 4.4% | 2 | Denmark, Europe |
| $\zeta_6$ | | ORF1ab | 21255 | G>C | | 80 | 4.4% | 1557 | USA, North America |
| $\zeta_7$ | | S | 21614 | C>U | L>F | 79 | 2.5% | 1442 | United Kingdom, Europe |

**Extended Data Table 2. SARS-CoV-2 variants in the 68KG dataset (continued).**
| Mutant<br>(major) | Mutant<br>(minor) | Gene | Genomic<br>Position | Nucleotide<br>change | Amino acid<br>change | Time<br>(days) | Variant<br>Frequency | Genomes<br>mapped |
| --- | --- | --- | --- | --- | --- | --- | --- | --- |
| $\varepsilon_1$ | | N | 28881 | G>A | R>K | 54 | 41.7% | 5 |
| $\varepsilon_2$ | | N | 28882 | G>A | R>K | 54 | 41.6% | 0 |
| $\varepsilon_3$ | | N | 28883 | G>C | G>R | 54 | 41.6% | 13394 |
| | $\varepsilon_{3a}$ | ORF1ab | 313 | C>U | | 64 | 2.4% | 1630 |
| | $\varepsilon_{3b}$ | ORF1ab | 19839 | U>C | | 64 | 2.9% | 1227 |
| | $\varepsilon_{3c}$ | M | 27046 | C>U | T>M | 69 | 0.8% | 548 |
| | $\varepsilon_{3d}$ | ORF1ab | 10097 | G>A | G>S | 69 | 3.2% | 11 |
| | $\varepsilon_{3e}$ | S | 23731 | C>U | | 69 | 3.2% | 425 |
| | $\varepsilon_{3f}$ | N | 28580 | G>U | D>Y | 69 | 1.0% | 678 |
| | $\varepsilon_{3g}$ | ORF1ab | 13536 | C>U | | 69 | 1.6% | 23 |
| | $\varepsilon_{3h}$ | ORF1ab | 4002 | C>U | T>I | 69 | 1.6% | 1066 |
| | $\varepsilon_{3i}$ | ORF1ab | 10265 | G>A | G>S | 63 | 1.4% | 879 |
| | $\varepsilon_{3j}$ | S | 21575 | C>U | L>F | 54 | 1.0% | 248 |
| | $\varepsilon_{3k}$ | S | 21637 | C>U | | 111 | 1.3% | 873 |
| | $\varepsilon_{3l}$ | ORF8 | 28169 | A>G | | 103 | 1.3% | 0 |
| | $\varepsilon_{3m}$ | ORF1ab | 16968 | G>U | | 114 | 1.0% | 702 |
| $\eta_1$ | | ORF1ab | 1163 | A>U | I>F | 86 | 9.6% | 339 |
| | $\eta_{1a}$ | ORF1ab | 14202 | G>U | | 159 | 1.1% | 7 |
| | $\eta_{1b}$ | ORF1ab | 19542 | G>U | M>I | 81 | 1.2% | 23 |
| | $\eta_{1c}$ | S | 22388 | C>U | | 90 | 1.2% | 21 |
| | $\eta_{1d}$ | N | 29466 | C>U | A>V | 91 | 1.2% | 4 |
| | $\eta_{1e}$ | ORF1ab | 19718 | C>U | T>I | 73 | 1.5% | 23 |
| | $\eta_{1f}$ | ORF3a | 26060 | C>U | T>I | 92 | 1.2% | 7 |
| | $\eta_{1g}$ | N | 29227 | G>U | | 55 | 1.2% | 24 |
| | $\eta_{1h}$ | ORF1ab | 3256 | U>C | | 167 | 1.1% | 0 |
| | $\eta_{1i}$ | ORF1ab | 5622 | C>U | P>L | 67 | 1.2% | 775 |
| $\eta_2$ | | ORF1ab | 18555 | C>U | | 51 | 8.0% | 25 |
| $\eta_3$ | | ORF1ab | 16647 | G>U | | 84 | 8.0% | 8 |
| $\eta_4$ | | ORF1ab | 7540 | U>C | | 86 | 7.9% | 0 |
| $\eta_5$ | | S | 23401 | G>A | | 86 | 7.9% | 1 |
| $\eta_6$ | | S | 22992 | G>A | S>N | 86 | 8.5% | 4583 |
| | $\eta_{6a}$ | S | 22480 | C>U | | 66 | 1.3% | 878 |
| $v_1$ | | ORF1ab | 17858 | A>G | Y>C | 59 | 2.6% | 61 |
| $v_2$ | | ORF1ab | 17747 | C>U | P>L | 59 | 2.5% | 1677 |

Note.- Genomic locations correspond to those of the NCBI genome (GenBank ID: NC\_04551.2). Amino acid changes are shown for nonsynonymous variants.

**Extended Data Figure 1.**
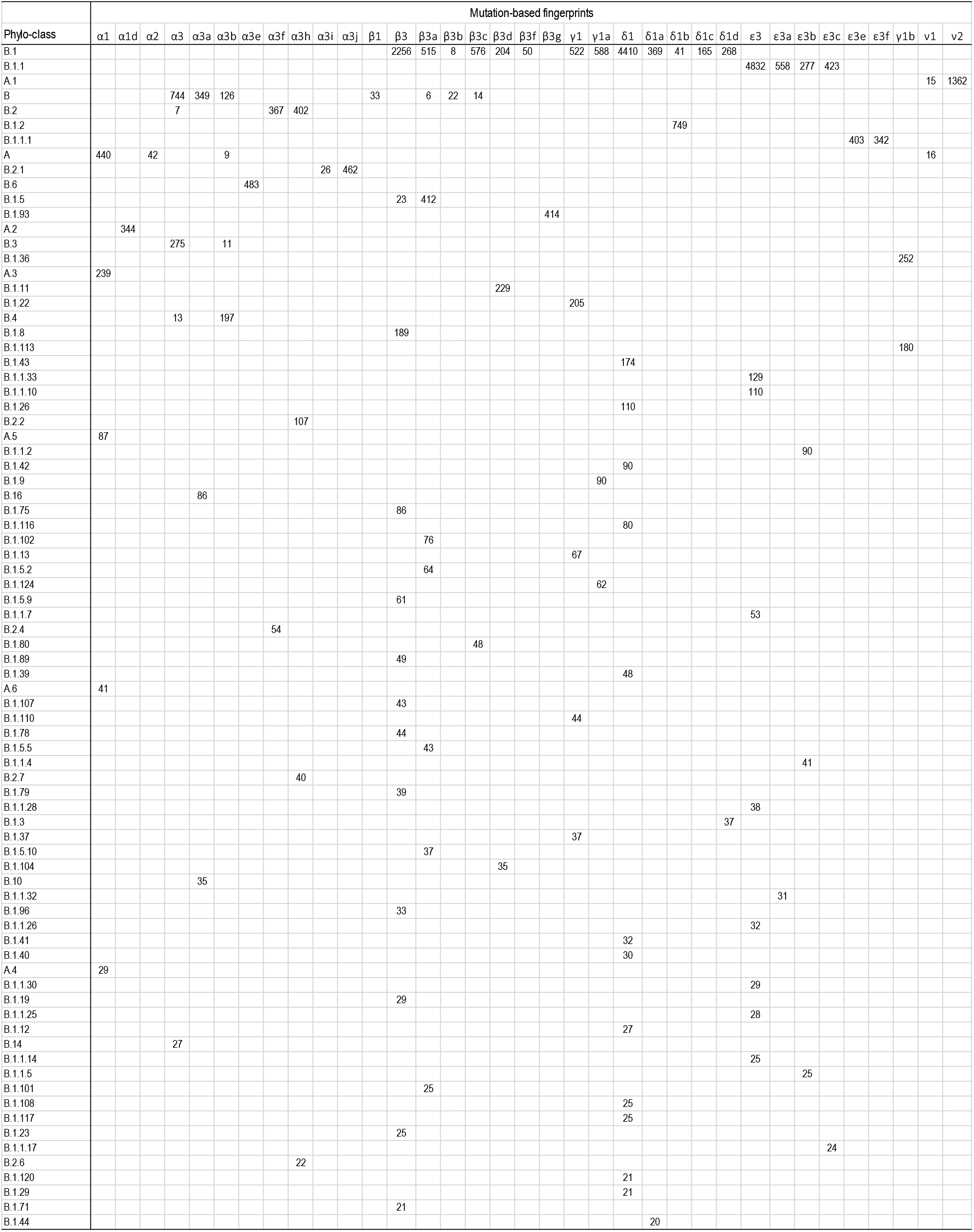
A comparison of mutation-based and phylogeny-based classifications of 29KG genomes. Phylogeny-based classification is obtained by using the Pangolin service (v2.0.3; https://pangolin.cog-uk.io/). Only the terminal variants are shown in mutation-based fingerprints for convenience. Each cell’s value is the number of genomes that belong to the corresponding mutation-based and phylogeny-based groups. All phylogenetic-based groups with fewer than 20 genomes are excluded. Cells with fewer than five genomes matching have been left empty to make the comparison more straightforward and allow for sequencing and estimation errors.

**Extended Data Figure 2.**
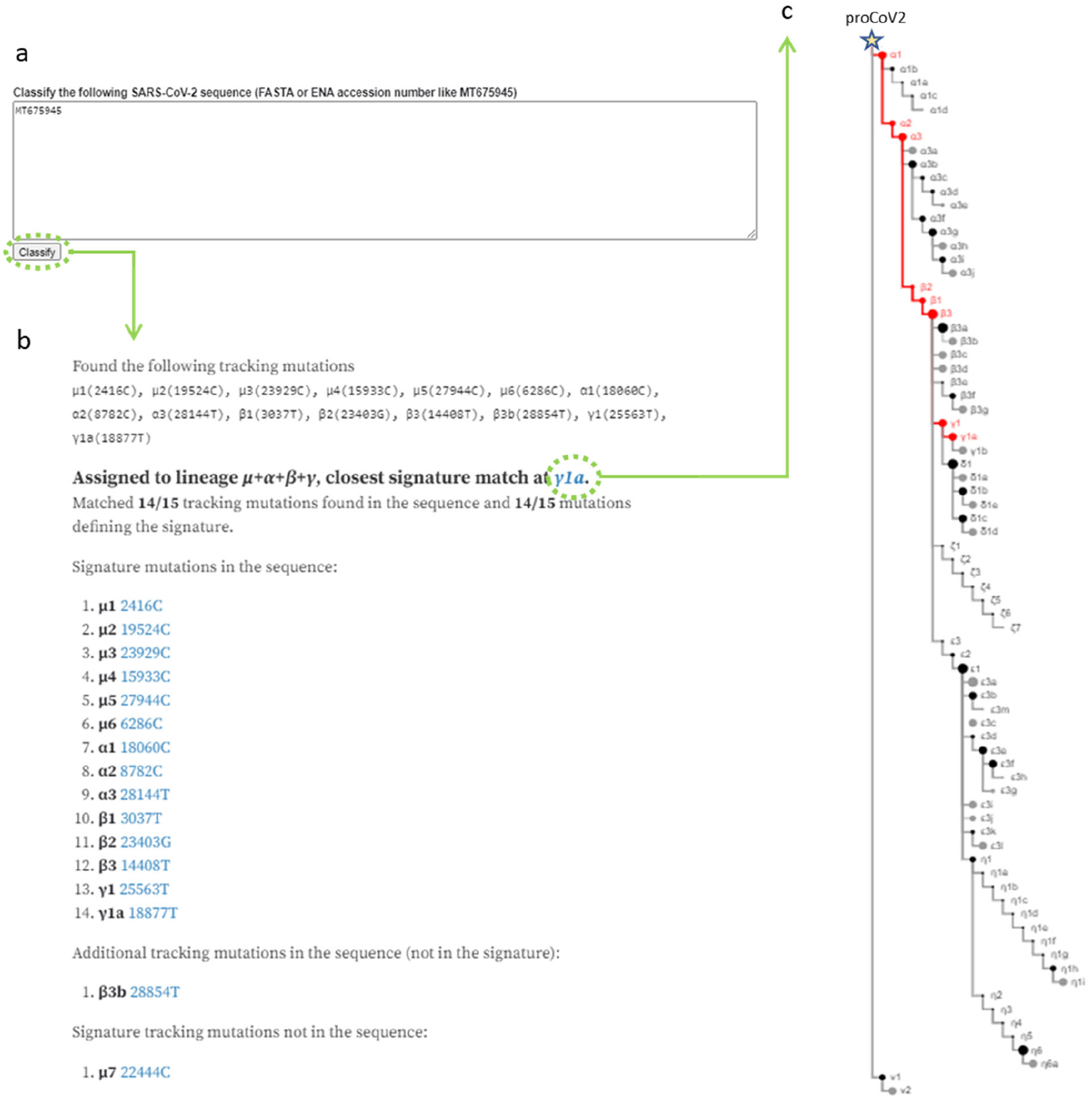
An example of sequence classification (ENA Accession MT675945) based on the 84 signature mutations (http://sars2evo.datamonkey.org/; “Classify your Sequence” option). (**a**) Input window to provide identifiers of sequences to be classified (e.g., MT675945). (**b**) The input sequence is classified into a mutational fingerprint. A list of mutations that are appeared in the input sequence is shown in the output window. (**c**) A waterfall phylogeny shows the input sequence’s location in the phylogeny, which appears after clicking the closet signature matched mutation in panel b.

